# High-throughput screening in hiPSC-cardiac models reveals cardiomyocyte-specific cell cycle regulatory mechanisms

**DOI:** 10.64898/2026.08.13.744744

**Authors:** Francesca Butera, Bryce Hassett, Rachel Morris, Jerico Revote, Hannah Huckstep, Lina H. H. Le, Jack Leerson, Thomas Martinez, Stephanie R. Hyslop, Sebastian Bass-Stringer, Antonia T. L. Zech, Tabitha Cree, Rebecca J. Sutton, Ivy K. N. Chiang, Eddy Kizana, Ellen B. Keen, James W. McNamara, Richard J. Mills, Sean J. Humphrey, Alejandro Hidalgo, Kevin I. Watt, David A. Elliott, Enzo R. Porrello

## Abstract

**Introductory Paragraph:** Multiple regulatory mechanisms govern cardiomyocyte proliferation including epigenetic modifications, metabolism and mechanical load. However, it is unclear whether such mechanisms can be pharmacologically targeted to induce cardiomyocyte proliferation without affecting other cell types. Here, we develop a dual-reporter (*TNNT2^eGFP^*; *PCNA^mScarlet-I^*) and a high-throughput image-based pipeline in human induced pluripotent stem cell (hiPSC)-derived cardiomyocytes, with counter screening in non-myocytes, to identify compounds that selectively promote cardiomyocyte proliferation without affecting other cell types. We identify the PIM kinase inhibitor GDC-0339 as a cardiomyocyte-selective pro-proliferative compound. GDC-0339 induced proliferation of hiPSC-derived cardiomyocytes without activity in non-myocytes, non-cardiac fibroblasts or epithelial cells. Phosphoproteomic profiling of GDC-0339 in cardiomyocytes and non-cardiac fibroblasts revealed a cardiomyocyte-specific mechanism of action involving sarcomere disassembly via remodelling of the F-actin cytoskeleton and metabolic reprogramming to anaerobic metabolism via Pyruvate Dehydrogenase Kinases (PDKs). Thus, we uncover cardiomyocyte-specific mechanisms governing the cell cycle that are potentially druggable.

## Main text

Fetal heart development relies on cardiomyocyte proliferation driven by signalling networks including WNT/μ-catenin (Lian et al., 2012; Quaife-Ryan et al., 2020), Hippo (Heallen et al., 2013), Notch (Campa et al., 2008), and neuregulin/ERBB (Bersell et al., 2009; D’Uva et al., 2015), which are postnatally suppressed in mature cardiomyocytes through epigenetic mechanisms (Butera et al., 2026). Furthermore, there is a metabolic switch at birth from glycolysis to fatty acid and oxidative metabolism that promotes cardiomyocyte cell cycle withdrawal and maturation (Puente et al., 2014; Mills et al., 2017; Cardoso et al., 2020; Li et al., 2023). In addition, sarcomere maturation and mechanical loading impose physical barriers to cardiomyocyte proliferation (Ahuja et al., 2004; Lam et al., 2025; Ciucci et al., 2026). While forced re-activation of proliferative transcriptional networks using genetic approaches can specifically activate the cell cycle in adult cardiomyocytes (Bersell et al., 2009; Xin et al., 2013; Leach et al., 2017; Gabisonia et al., 2019; Liu et al., 2021; Li et al., 2023), it is unclear whether pharmacological approaches could be developed to specifically promote cardiomyocyte proliferation without affecting other cell types. In this regard, induced pluripotent stem cells provide a useful high-throughput human model system to screen for small molecules that solely promote proliferation of differentiated cardiomyocytes (Mills et al., 2019).

We performed an initial screen of 775 compounds targeting diverse signalling pathways including cell cycle regulators and epigenetic modifiers to identify cardiomyocyte-selective pro-proliferative compounds using fixed hiPSC-cardiac monolayers (Fig. 1a). To find molecules effective in both healthy and diseased cardiomyocytes, we used hiPSCs derived from a healthy donor (PB10.5), as well as an isogenic pair of hiPSCs varying only at the pathogenic mutation c.4246_4247del (p.Leu1416AsnfsTer23) in Desmoplakin (denoted MCHTB11.9; hereafter *DSP^mut^* and H3; hereafter *DSP^corr^*) (Pocock et al., 2025) (Extended Data Fig. 1a,b). Compound 6.28 (C6.28), a pro-proliferative molecule that inhibits both GSK3 and MST1, was included as a positive control (Mills et al., 2017, 2019) (Fig. 1a). In total, 39 compounds were defined as pro-proliferative (% of pRB+/post-G1 cardiomyocytes z-score > 1.75; in 2 of 3 lines tested) and non-toxic in cardiomyocytes (Extended Data Fig. 1c-e and Extended Data Table 1).

**Fig. 1.**
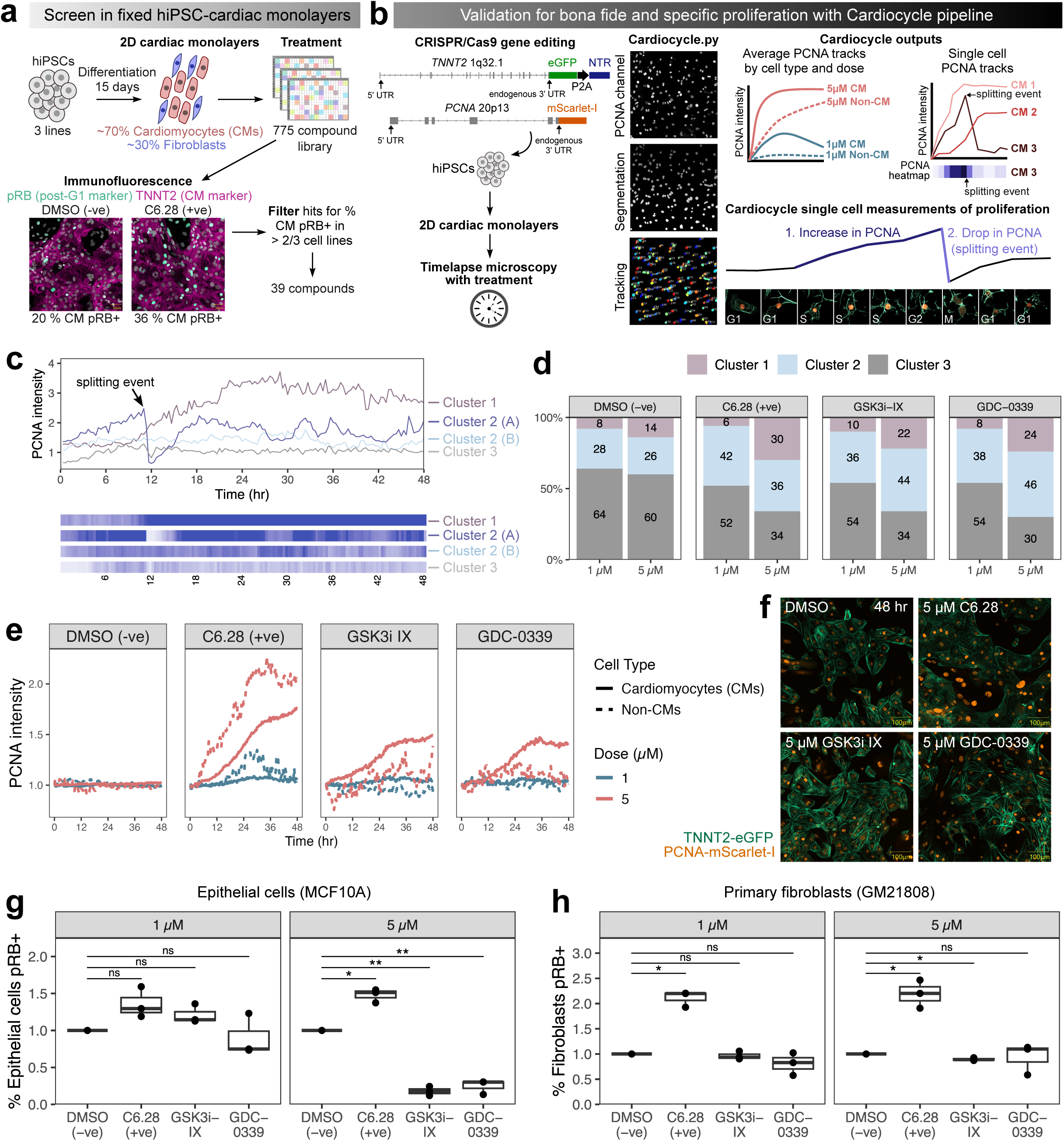
Cardiocycle pipeline identifies GSK3i-IX and GDC-0339 as compounds enhancing cardiomyocyte-specific proliferation. a. Schematic of high-throughput screen of 775 compounds in fixed hiPSC-cardiac monolayers. Following 48 hr compound treatment, hiPSC-cardiac cells were fixed and stained for pRB to mark post-G1 cells and TNNT2 to mark cardiomyocytes. Example images are shown from DSP^corr^ (H3) hiPSC-cardiac cells. Compounds were z-score ranked for the percentage of cardiomyocytes positive for pRB for each cell line and dose. The 775 compounds were filtered to 39 based on a z-score > 1.75 for either dose (1 or 5 µM) in two or more cell lines b. A novel workflow called Cardiocycle was developed for tracking cardiac cell proliferation in cardiomyocytes and non-cardiomyocytes in hiPSC-derived cultures imaged by timelapse microscopy. This workflow combines a novel automated image analysis (Cardiocycle.py) with a dual reporter hiPSC line generated by CRISPR/Cas9 gene editing expressing cTnT-eGFP and Proliferating Cell Nuclear Antigen (PCNA)-mScarlet-I tagged at the endogenous loci. Cardiocycle assesses proliferation two-fold: 1) an overall increase in PCNA intensity and 2) a drop in PCNA intensity indicating a cytokinetic/’splitting’ event. The pipeline outputs single cell PCNA tracks as well as average tracks per condition (cell type and treatment). c. Representative single nucleus tracks of PCNA intensity from the three hierarchical clusters. Cluster 1 contains nuclei with have sustained high PCNA. Cluster 2 contains nuclei with moderate PCNA and includes tracks where a splitting event occurs (e.g. track labelled ‘Cluster 2A’), whereby PCNA intensity suddenly drops, marking transition from G2 to the daughter cell G1. Cluster 3 contains nuclei with sustained low PCNA, indicating a lack of cell cycle activity. d. Proportion of hiPSC-cardiomyocytes in each PCNA cluster for each compound treatment and dose (1 µM or 5 µM), showing that GSK3i-IX and GDC-0339 increase the proportion in the cluster associated with cell cycle activity and division (Cluster 2). e. Tracks of mean PCNA-mScarlet-I intensity (by cell type and dose) over time for hiPSC-cardiac cells treated with DMSO (negative control), Compound 6.28 (positive control), GSK3i-IX and GDC-0339 showing cardiomyocyte-specific upregulation of PCNA by GSK3i-IX and GDC-0339. PCNA-mScarlet-I intensity was quantified by living imaging of hiPSC-cardiomyocyte monolayers at 20 min intervals. Cardiomyocyte/TNNT2+ cells are represented by a solid line and non-cardiomyocyte/TNNT2-cells are represented by a dashed line. f. Representative images of hiPSC-cardiac monolayers expressing PCNA-mScarlet-I (orange) and TNNT2-eGFP (green) treated with compounds for 48 hr. g. Percentage of MCF10A breast epithelial cells positive for the post-G1 marker phospho-RB. Cells were treated with DMSO (negative control), Compound 6.28 (positive control), GSK3i-IX or GDC-0339 for 48 hr at 1 or 5 µM. h. Percentage of GM21808 primary fibroblasts positive for the post-G1 marker phospho-RB. Cells were treated with DMSO (negative control), Compound 6.28 (positive control), GSK3i-IX or GDC-0339 for 48 hr at 1 or 5 µM.

Fixed immunofluorescence assays cannot distinguish between post-G1 cell cycle arrest versus cell proliferation, since cell populations in both states are positive for post-G1 markers (including pRB, Ki67 and PHH3) (Gerdes et al., 1984; Buchkovich et al., 1989; Hendzel et al., 1997). Therefore, a hiPSC reporter line was generated expressing eGFP from the *TNNT2* locus (*TNNT2^eGFP^*) and a PCNA:mScarlet-I fusion protein from the *Proliferating Cell Nuclear Antigen* locus (*PCNA^mScarlet-I^*; Fig. 1b and Supplementary Movie 1). For analysis of PCNA:mScarlet-I changes over time, we developed Cardiocycle.py, a Python-based image analysis workflow that segments and tracks single nuclei to provide quantification of temporal PCNA changes (Fig. 1b and Extended Data Fig. 2a) (Butera et al., 2024). Cardiocycle.py reads out bona fide proliferation by detecting: 1) increases in PCNA intensity, which correlates with cell cycle progression as PCNA is required for DNA replication (Prelich et al., 1987) and 2) a drop in PCNA intensity indicating a cytokinetic/‘splitting’ event (Fig. 1b and Supplementary Movie 1). Profiling the 39 hit compounds from the static screen using our live imaging format identified three distinct PCNA responses within a data set of 228,896 cardiomyocyte nuclei tracks and 19,165 non-cardiomyocyte nuclei tracks (n = 3, at 1 µM or 5 µM for 48 hr); namely, continued increased intensity (Cluster 1), those with a moderate increase in intensity and identifiable cell division events (Cluster 2a,b) and no or little change (Cluster 3) (Fig. 1c and Extended Data Fig. 2b). Focusing on *TNNT2^eGFP+ve^* cardiomyocytes, the GSK3 inhibitor, GSK3i-IX, and the PIM Kinase inhibitor, GDC-0339, were shown to be cardiomyocyte-specific pro-proliferative molecules (Fig. 1d-f and Extended Data Fig. 2c-e). GSK3i-IX (also called BIO) induces proliferation of adult rat cardiomyocytes, supporting the validity of our pipeline (Tseng et al., 2006). Other compounds induced cardiomyocyte proliferation but also increased non-cardiomyocyte proliferation (e.g. At9283, Jnj-7706621 and Unc1999), caused only mild cardiomyocyte-specific PCNA increases (e.g. Atuveciclib) or were autofluorescent false-positives (e.g. Proflavine) (Extended Data Fig. 2c). Upregulation of pRB in the primary screen by the remaining compounds may reflect induction of S or G2 phase cell cycle arrest. Neither GSK3i-IX or GDC-0339 promoted proliferation in an epithelial cell line (MCF10A) (Fig. 1g) or primary non-cardiac fibroblasts (Fig. 1h), suggesting these compounds may act in a cardiomyocyte-specific manner. In contrast, the mitogen C6.28 increased the percentage of pRB+ proliferative cells in both cell types (Fig. 1g,h).

We next tested whether GSK3i-IX and GDC-0339 induced cardiomyocyte cell cycle activity in more mature three-dimensional hiPSC-cardiac organoids from a PB10.5 background (Pocock et al., 2025) (Fig. 2). After 48 hours of GDC-0339 exposure, 30 % of cardiomyocytes expressed Ki67 across three doses (2.5, 5 or 10 µM). Similarly, GSK3i-IX induced Ki67 in almost 50 % of cardiomyocytes at low doses, while at 10 µM there was no difference with the DMSO control (Fig. 2a,b). To confirm that cardiomyocytes were entering the cell cycle, the CDK4/6 inhibitor Palbociclib, which prevents passage through the G1/S cell cycle checkpoint (Fry et al., 2004), was used. Palbociclib alone effectively reduced Ki67 incidence from 18 % of cardiomyocytes in the DMSO control to 4 % at 150 nM and to 3 % at 1 µM Palbociclib (Fig. 2c, d). In the presence of GSK3i-IX (5 µM) and GDC-0339 (5 µM), 150 nM Palbociclib halved Ki67+ incidence and at 1 µM reduced Ki67+ incidence to < 3 %. Consistent with this finding, pRB levels were dramatically reduced in the presence of Palbociclib (Fig. 2e, f). These data were replicated in cardiac organoids generated from an independent embryonic stem cell line (*NKX2-5^eGFP/w^*; Elliott et al., 2011) (Extended Data Fig. 3). Collectively, these data demonstrate that GSK3i-IX and GDC-0339 can drive more mature hiPSC-derived cardiomyocytes into cycle.

**Fig. 2.**
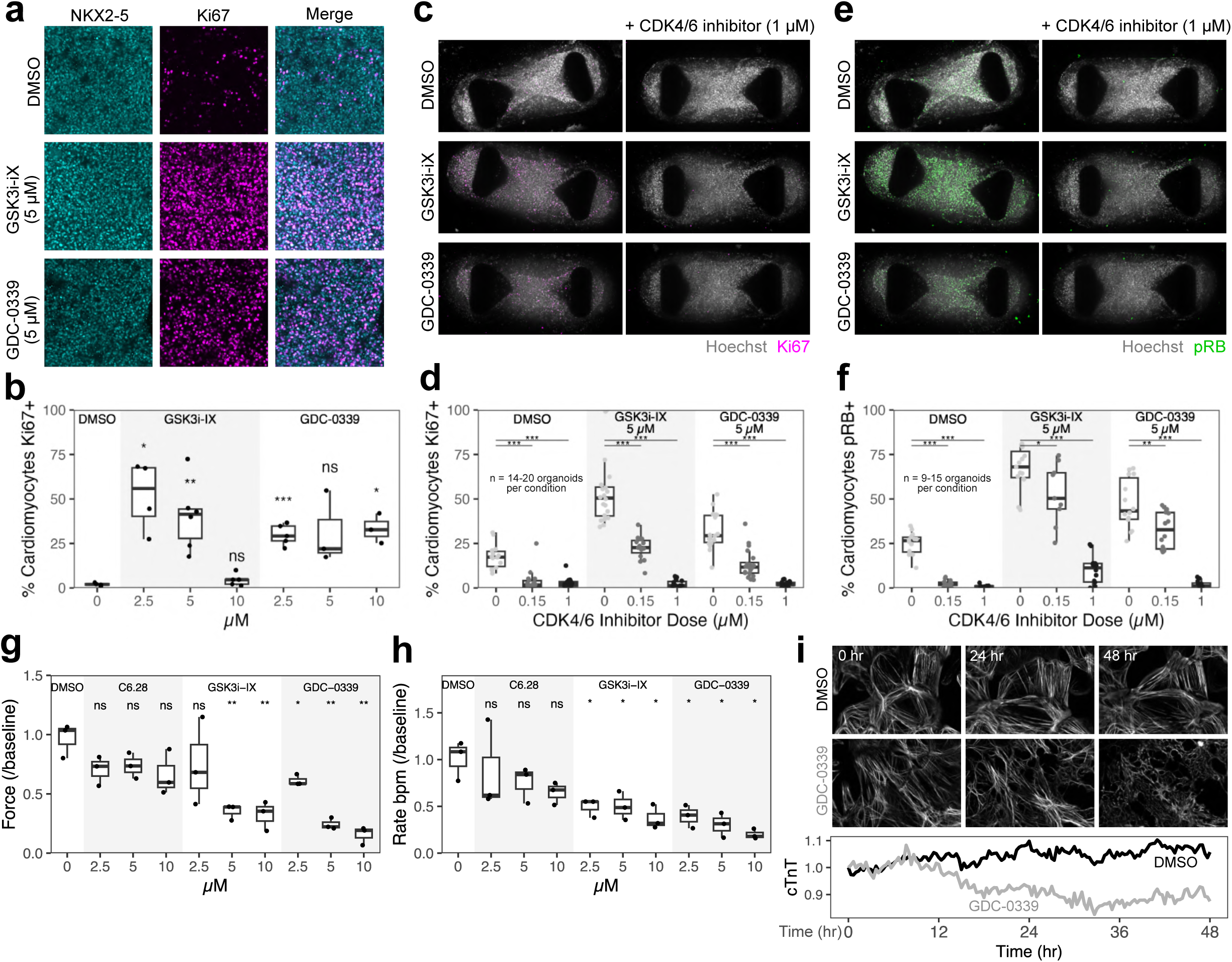
GSK3i-IX and GDC-0339 enhance cell cycle activation in hiPSC-cardiac organoids while impairing contraction. a. Representative images of NKX2-5 and Ki67 from immunofluorescence staining of hiPSC-cardiac organoids treated with DMSO, GSK3i-IX (2.5, 5 and 10 µM) and GDC-0339 (2.5, 5 and 10 µM) for 48 hr. The center field per organoid is shown. b. Percentage of cardiomyocytes positive for the cell cycle marker Ki67 in hiPSC-cardiac organoids from wildtype PB10.5 background. Organoids were treated with DMSO, GSK3i-IX (2.5, 5 and 10 µM) and GDC-0339 (2.5, 5 and 10 µM) for 48 hr. NKX2-5 was used to distinguish cardiomyocytes from non-cardiomyocytes. Each point represents an organoid. Data shown from 1 experiment. c. Representative images of Hoechst and Ki67 staining in hiPSC-cardiac organoids treated for 48 hr with DMSO, GSK3i-IX (5 µM) or GDC-0339 (5 µM) for 48 hr with co-treatment with the CDK4/6 inhibitor Palbociclib at 0.15 µM or 1 µM. d. Percentage of cardiomyocytes positive for the cell cycle marker Ki67 in hiPSC-cardiac organoids from PB10.5 ACTN2-Scarlet hiPSCs. Organoids were treated for 48 hr with DMSO, GSK3i-IX (5 µM) or GDC-0339 (5 µM) for 48 hr with co-treatment with the CDK4/6 inhibitor Palbociclib at 0.15 µM or 1 µM. ACTN2-Scarlet (not shown) was used to distinguish cardiomyocytes from non-cardiomyocytes. Each point represents an organoid (14-20 per condition). Data shown from 3 biological replicates. e. Representative images of Hoechst and pRB staining in hiPSC-cardiac organoids treated for 48 hr with DMSO, GSK3i-IX (5 µM) or GDC-0339 (5 µM) for 48 hr with co-treatment with the CDK4/6 inhibitor Palbociclib at 0.15 µM or 1 µM. f. Percentage of cardiomyocytes positive for the cell cycle marker pRB in hiPSC-cardiac organoids from PB10.5 ACTN2-Scarlet hiPSCs. Organoids were treated for 48 hr with DMSO, GSK3i-IX (5 µM) or GDC-0339 (5 µM) for 48 hr with co-treatment with the CDK4/6 inhibitor Palbociclib at 0.15 µM or 1 µM. ACTN2-Scarlet (not shown) was used to distinguish cardiomyocytes from non-cardiomyocytes. Each point represents an organoid (9-15 per condition). Data shown from 3 biological replicates. g. Contractility (pole tracking) analysis of force in hiPSC-cardiac organoids from PB10.5 background. Organoids were treated with compounds in maturation media for 48 hr from day 22 of the differentiation protocol and fixed on day 24. Measurements from 24 are normalised to the measurement on day 22 by each organoid. Points represent experimental replicates (n = 3), calculated as the mean of 2-7 organoids per experiment and condition. h. Contractility analysis of beat rate. i. Quantification of mean intensity and representative images of TNNT2-eGFP from 48 hr timelapse microscopy with 20 min intervals. PB10.5 hiPSC-cardiac monolayers were treated with DMSO or GDC-0339 (5 µM). Data are normalised to the DMSO measurement at the first timepoint.

Sarcomere disassembly is a prerequisite for cardiomyocyte proliferation, therefore pro-proliferative compounds may impair contractility (Ahuja et al., 2004; Mills et al., 2019; Morikawa et al., 2025). Both GSK3i-IX and GDC-0339 dramatically reduced force generation in hiPSC-cardiac organoids (Fig. 2g). Further, the beat rate of cardiac organoids treated with GSK3i-IX and GDC-0339 was significantly lower (Fig. 2h). Consistent with the reduction in force, *TNNT2^eGFP^*;*PCNA^mScarlet-l^* cardiac monolayers treated with GDC-0339 had significantly lower levels of *TNNT2:eGFP* fluorescence by 48 hours (Fig. 2i). Taken together, these data show that sarcomere integrity and cardiac contractility are impaired during induction of cardiomyocyte proliferation by GSK3i-IX and GDC-0339.

To identify core initiating signalling events required for cardiomyocyte proliferation, we performed deep phosphoproteomics to define the phosphosites that respond rapidly and selectively in hiPSC-derived cardiac cells in response to mitogen treatment (Fig. 3a), using the kinase inhibitors GDC-0339, GSK3i-IX and C6.28 across a 7-point dose range (0, 0.375, 0.75, 1.5, 3, 5 and 10 µM). To attain cardiomyocyte-specific phosphosites regulated during proliferation, independent treatment of primary non-cardiac fibroblasts was included. In total, over 140,000 phosphopeptides were analysed (Fig. 3a). A subset of 376 phosphosites were differentially regulated exclusively in cardiomyocytes with at least two of the three compounds in the same direction (Extended Data Table 2). The top cardiomyocyte-specific phosphosites overlapping between the proliferative compounds were in proteins linked to mechanotransduction and cytoskeletal remodelling, including DSP, KANK2, SVIL, MAP1A, CTNND1 and ARVCF (Fig. 3b). Kinase substrate enrichment analysis (KSEA) revealed upregulation of several proliferation-associated kinases, including CDK1, CDK2, MAPK1 and MAPK3 (Fig. 3b). In addition, mitogen treatment upregulated the metabolic kinases Pyruvate Dehydrogenase Kinase 1-3 (PDK1-3) and the cytoskeleton regulator P21-Activated Kinase 1 (PAK1), which is implicated in regulating cardiac contractility (Sheehan et al., 2007). Corresponding gene set enrichment analysis (GSEA) revealed a cardiomyocyte-specific phosphosignature positively enriched for the Gene Ontology (GO) Biological Process ‘mRNA processing’ (Fig. 3c), indicative of increased transcription, and downregulation of the term ‘actin-filament-based process’, suggesting remodelling of the actin cytoskeleton. Clustering of the 115 cardiomyocyte-specific mitogen-regulated phosphosites (corresponding to 55 genes) associated with ‘actin-filament-based process’ revealed that phosphosites upregulated with the initiation of cardiomyocyte proliferation have a binary response observed at the lowest mitogen concentration 0.375 µM (Fig. 3d). In contrast, downregulated phosphosites display a more dose-dependent response, with the majority of phosphosites downregulated upwards of 3 µM.

**Fig. 3.**
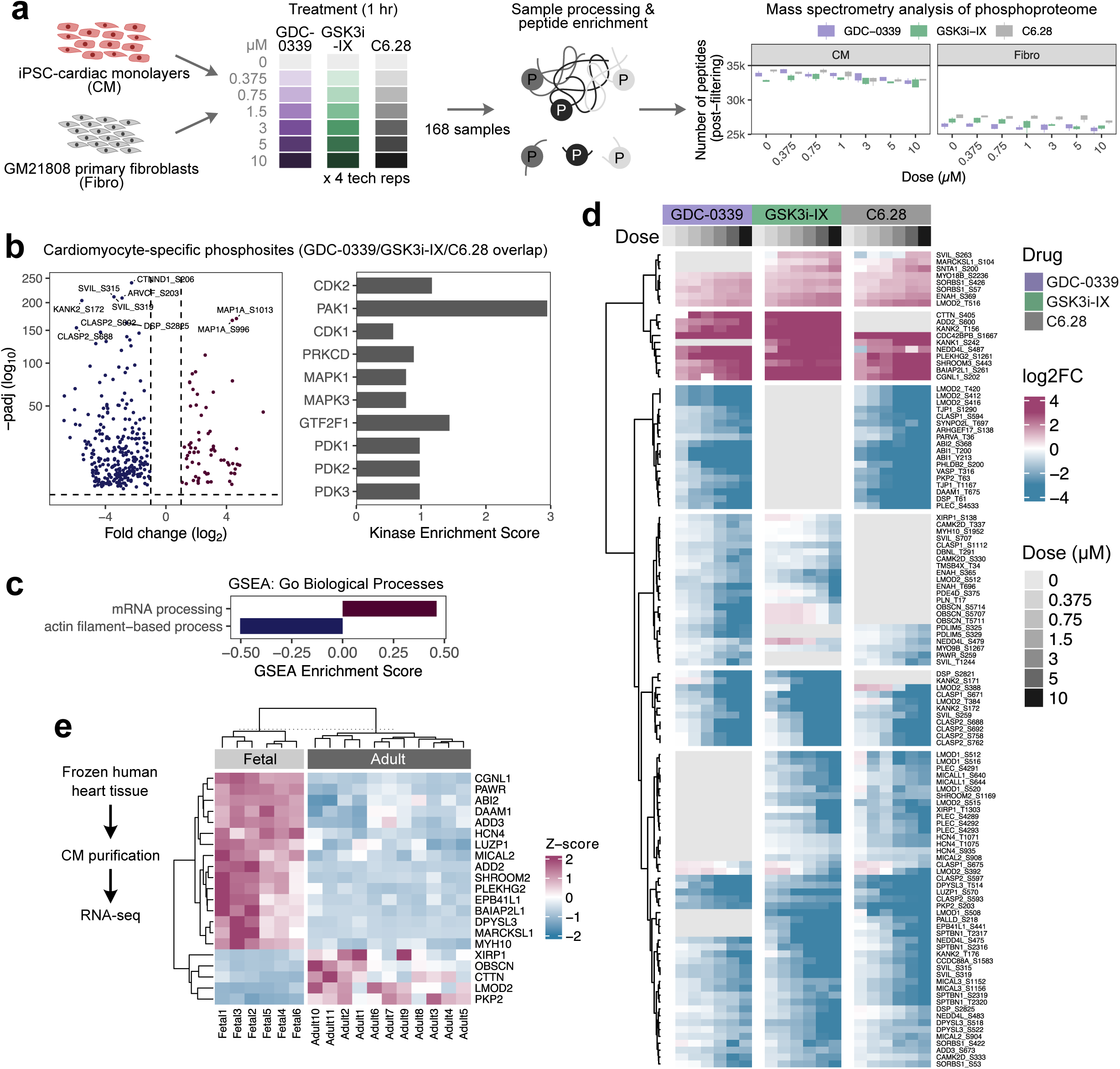
Phosphoproteomics reveals that actin remodelling is a core process associated with cardiomyocyte proliferation. a. Schematic overview of phosphoproteomics experiment. HiPSC-cardiomyocytes (CM) and non-cardiac GM21808 primary fibroblasts (Fibro) were processed separately. Cells were treated for 1 hr with GSK3i-IX, GDC-0339 or Compound 6.28 (C6.28) at a 7-point dose range (0, 0.375, 0.75, 1.5, 3, 5 and 10 µM), with four biological replicates per condition. Following sample lysis, peptides were enriched then detected using the Orbitrap Astral mass spectrometer. Post-filtering, 30,000-35,000 peptides were detected for each cardiomyocyte sample and 25,000-30,000 peptides were detected for each fibroblast sample. b. Left: Volcano plot for overlapping differentially regulated phosphosites with at least two of the three cardiomyocyte mitogens GSK3i-IX, GDC-0339 or C6.28 in the same direction and exclusively in cardiomyocytes Cardiomyocyte-specificity was attained by filtering out phosphosites differentially regulated in fibroblasts. Differentially regulated phosphosites were determined using the DESeq2 Likelihood Ratio Test across 7-point dose curve independently by compound and cell type. Right: Top 10 kinases ranked by pvalue (most significant at top) from Kinase Substrate Enrichment Analysis (KSEA) using the overlapping cardiomyocyte-specific phosphosites. c. Gene Set Enrichment Analysis (GSEA) for differentially regulated cardiomyocyte-specific phosphosites overlapping with GSK3i-IX, GDC-0339 or C6.28 treatment (2/3 compounds in the same direction). d. Clustered heatmap of the overlapping cardiomyocyte-specific phosphosites in hiPSC-cardiac cells associated with the GO biological process ‘actin-filament based process’ (115 phosphosites corresponding to 55 genes). Each heatmap column represents the mean of four biological replicates for each compound and dose. The heatmap is coloured by log2 fold-change to the respective DMSO control for each compound. e. Heatmap showing gene expression for the 21 out of 55 genes associated with the 115 cardiomyocyte-specific phosphosites in (d) with statistically significant expression in cardiomyocytes from primary human fetal versus adult heart tissue (t-test with Benjamini-Hochberg multiple test correction; padj < 0.001). Data originally published by Sim et al. (2021). Bulk RNA-seq was performed on nuclei isolated from human hearts stratified by developmental stage. Fetal samples from 14-20 weeks gestation (n = 6 individuals) and adult samples from 35-65 years age (n = 11 individuals). Samples are clustered following stratification by developmental stage.

Consistent with the notion that actin dynamics play a critical role in cardiomyocyte proliferation, 21/55 genes encoding the actin-associated cardiomyocyte-specific phosphosites (Fig. 3d) displayed reciprocal transcriptional expression profiles between fetal and adult cardiomyocytes from primary human heart tissue (padj < 0.001 for t-test with Benjamini-Hochberg multiple test correction) (Fig. 3e) (Mehdiabadi et al., 2022; Sim et al., 2021). 5 genes are upregulated in adult cardiomyocytes: XIRP1, OBSCN, CTTN, LMO2 and PKP2 (Fig. 3e). Meanwhile, 16 genes are downregulated in adult cardiomyocytes, including the actin-capping protein adducin 2 (*ADD2*), which has been recently characterised as a regulator of sarcomere disassembly (Xiao et al., 2024), a prerequisite for cardiomyocyte proliferation (Pettinato et al., 2022). Thus, the phosphorylation of these components of the actin-cytoskeletal network appears to be an important initiating step in the cardiomyocyte cell cycle.

Given that the phosphosignature of hiPSC-cardiomyocytes induced to proliferate was dominated by actin signalling (Fig. 3c-d), we quantified filamentous actin (F-actin) morphology in hiPSC-cardiac monolayers by Phalloidin staining (Extended Data Fig. 4). Overall changes in F-actin abundance (Intensity), changes in F-actin abundance relative to background (Contrast), heterogeneity in abundance across the field of view (Variance), and regularity of F-actin across the field of view (Angular Second Moment/ASM and Entropy) were obtained (Extended Data Fig. 4a) (Haralick et al., 1973). hiPSC-cardiac cells treated with proliferative compounds demonstrated disrupted F-actin morphology compared to the DMSO control (Extended Data Fig. 4b). While GSK3i-IX, GDC-0339 and C6.28 induced distinct F-actin morphology, actin remodelling was a clear common feature (Extended Data Fig. 4b-f). C6.28 resulted in more uniform F-actin intensity (low contrast and variance) with increased intensity at 5 µM, but minimal effect on F-actin distribution (Extended Data Fig. 4d). In contrast, GSK3i-IX and GDC-0339 reduced overall F-actin intensity while altering F-actin organisation (Extended Data Fig. 4e and Extended Data Fig. 4f). Further, GSK3i-IX caused an alteration in broad F-actin structure (high ASM and low entropy; Extended Data Fig. 4e) while GDC-0339 treatment led to smaller/more local changes in F-actin morphology, with reduced correlation between parallel F-actin filaments (Extended Data Fig. 4f). In contrast, MCF10A breast epithelial cells (Extended Data Fig. 5a-e) and primary dermal fibroblasts (Extended Data Fig. 5f-j) had milder disruption of F-actin morphology with significant overlap with DMSO controls (Extended Data Fig. 5a, f). Taken together, these findings suggest that cardiomyocyte proliferation involves substantial actin remodelling in response to cardiomyocyte mitogens, which does not occur to the same extent in non-myocytes.

Next, we focused on elucidating the pathways modulated by GSK3i-IX and GDC-0339 that may mediate cardiomyocyte-specific cell cycle re-entry. KSEA with differentially regulated phosphosites from acute GSK3i-IX treatment in hiPSC-cardiac cells identified downregulation of the expected kinases GSK3α/β (Extended Data Fig. 6a). Comparing differentially regulated phosphosites by GSK3i-IX in hiPSC-cardiac monolayers to dermal fibroblasts displayed highly cell type-specific phosphosignatures. Overall, 29.1 % of identified phosphosites were shared between cardiomyocytes and dermal fibroblasts, 13.7 % were unique to cardiomyocytes and 57.2 % were unique to fibroblasts (Extended Data Fig. 6b; Extended Data Table 3).

Consistent with the known role of GSK3α/β-inhibition in disrupting contractility (Zhou et al., 2016; Gupte et al., 2018; Mills et al., 2019), many of the top cardiomyocyte-specific phosphosite changes induced by GSK3i-IX (261 phosphosites increased; 212 reduced; |log2foldchange| > 1; padj < 0.05) were found in cytoskeletal or contractile proteins, such as SVIL, RCSD1 and LMOD2 (Extended Data Fig. 6c). Focusing on the cardiomyocyte-specific phosphosignature, KSEA again identified GSK3A and GSK3B, suggesting these kinases phosphorylate some targets exclusively in cardiomyocytes (Extended Data Fig. 6d). Other kinases impacted by GSK3i-IX include the cell cycle regulators MAPK1 (also known as ERK1) and AKT1.

We then focused on the cardiomyocyte-specific mitogen GDC-0339. KSEA with GDC-0339-regulated phosphosites indicated inhibition of the compound’s known targets PIM1-3 and GSK3B (Fig. 4a) (Wang et al., 2019), in addition to inhibition of the ribosomal kinases RPS6KB1/2 and activation of PDK1-3. Similar to GSK3i-IX, analysis of GDC-0339-regulated phosphosites revealed highly cell type-specific phosphorylation signatures, with only 30.9 % of GDC-0339-regulated phosphosites shared by cardiomyocytes and fibroblasts (Fig. 4b; Extended Data Table 4). In cardiomyocytes specifically, GDC-0339 upregulated 291 phosphosites and downregulated 161 phosphosites (|log2foldchange| > 1; padj < 0.05) (Fig. 4c). Surprisingly, KSEA with cardiomyocyte-specific phosphosites revealed that GDC-0339 is predicted to have a more activatory function in cardiomyocytes, including activation of all four PDKs and the cell growth-associated kinase JAK2 (Fig. 4c).

**Fig. 4.**
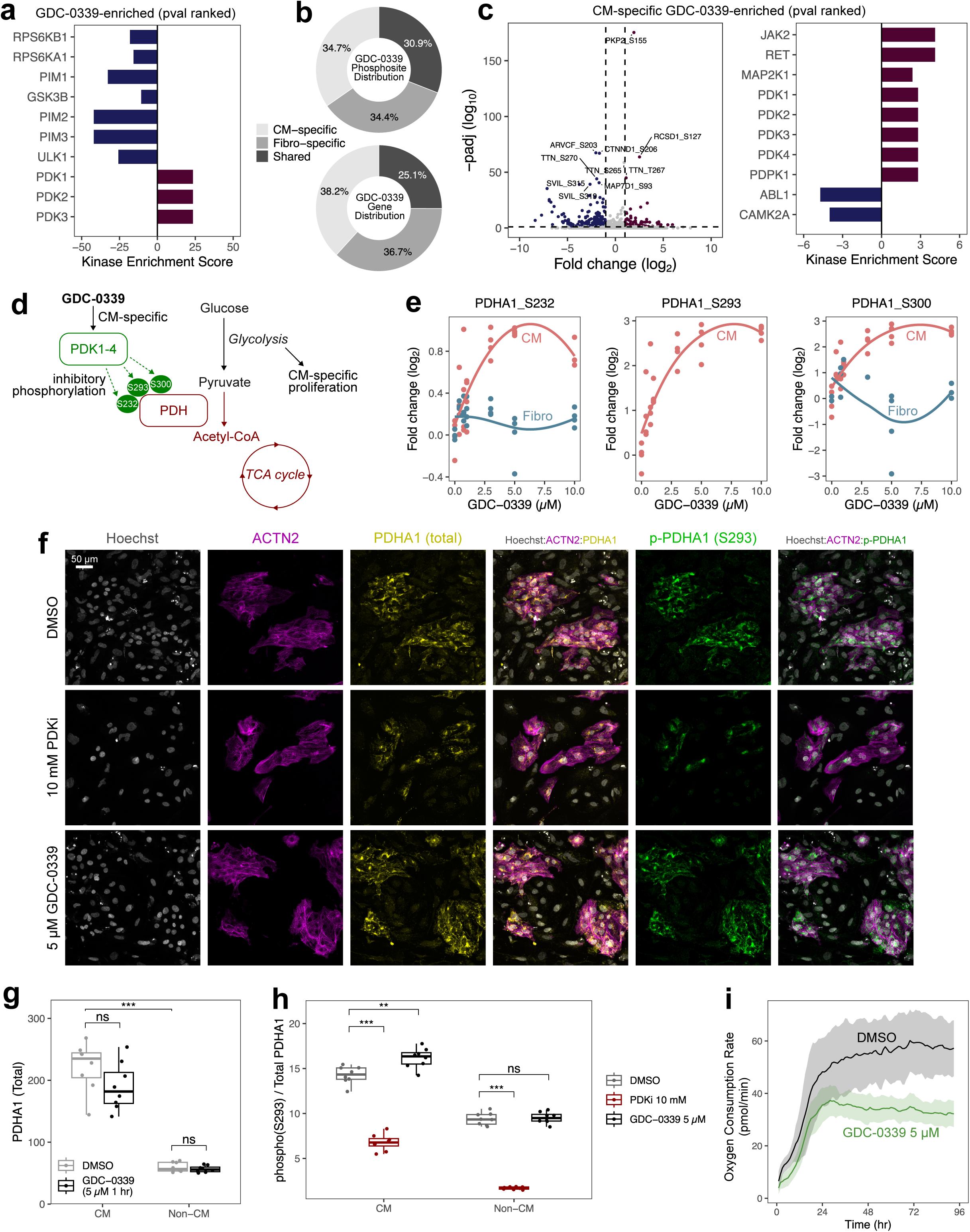
GDC-0339 rewires metabolism in cardiomyocytes by upregulating PDHA1 phosphorylation and reducing oxygen consumption. a. Kinases identified by KSEA for differentially-regulated phosphosites by GDC-0339 in hiPSC-cardiac cells (irrespective of effect in dermal fibroblasts). Upregulated kinases (positive enrichment score) in red, downregulated kinases (negative enrichment score) in blue. b. Top: percentage of differentially regulated phosphosites by GDC-0339 classed as statistically significant (DESeq2 Likelihood Ratio Test padj < 0.05, |log2foldchange| > 1) for cardiomyocytes alone (CM-specific), fibroblasts alone (Fibro-specific), or in both cell types (shared). Bottom: percentage of cardiomyocyte-specific, fibroblast-specific, or shared genes associated with the differentially regulated phosphosites by GDC-0339. c. Left: Volcano plot for cardiomyocyte-specific differentially regulated phosphosites by GDC-0339 (DESeq2 Likelihood Ratio Test padj < 0.05, |log2foldchange| > 1). Right: Top 10 kinases ranked by pvalue (most significant at top) from Kinase Substrate Enrichment Analysis (KSEA) using the GDC-0339-regulated cardiomyocyte-specific phosphosites. d. Schematic of proposed signalling mechanism for GDC-0339 induction of cardiomyocyte-specific proliferation. Phosphoproteomics indicated significant enrichment of PDK activity selectively in cardiomyocytes. PDKs function by negative regulation of the PDH complex by phosphorylation of the PDHA1 component at the residues S232, S293 and S300. The PDH complex is required for conversion of pyruvate from glycolysis to acetyl-CoA, which enters the TCA cycle as a part of oxidative metabolism. Blocking PDHA1 increases reliance on glycolysis, a metabolic process associated with cardiomyocyte proliferation. e. Quantification by phosphoproteomics and mass spectrometry of PDHA1 residues phosphorylated by PDKs (S232, S293 and S300) for increasing GDC-0339 dose (0, 0.375, 0.75, 1.5, 3, 5 and 10 µM). Data shown for cardiomyocytes (CM; coral) and non-cardiac fibroblasts (Fibro; blue). f. Representative images of ACTN2-Scarlet and Hoechst, PDHA1 and phospho-PDHA1 (S293) staining in hiPSC-cardiac monolayers with 1 hr DMSO, 10 mM PDKi (DCA), or 5 µM GDC-0339 treatment. Images demonstrate cardiomyocyte-localised expression of PDHA1 and phospho-PDHA1. g. PDHA1 intensity in cardiomyocytes and non-cardiomyocytes (cardiac fibroblasts) treated with DMSO or GDC-0339 (5 µM) for 1 hr. h. phospho-PDHA1 (S293) intensity relative to total PDHA1 intensity calculated on a single cell level and shown as well means for cardiomyocytes and non-cardiomyocytes (cardiac fibroblasts). HiPSC-cardiac monolayers were treated for 1 hr with DMSO, GDC-0339 (5 µM) or PDKi (10 mM). i. Oxygen consumption rate over 96 hr measured in hiPSC-cardiac monolayers treated with DMSO or GDC-0339 (5 µM). n = 16 technical replicates per condition.

PDK1-4 mediate a switch in substrate utilisation between glycolysis and oxidative metabolism by phosphorylation of the PDH complex (Kolobova et al., 2001), which regulates the conversion of glycolytic pyruvate to acetyl-CoA that enters the TCA cycle (Patel and Roche, 1990). Canonically, PDK phosphorylation of the PDH component PDHA1 at the three specific residues S232, S293, and S300 negatively regulates PDH activity (Fig. 4d). In hiPSC-cardiac monolayers, GDC-0339 upregulated phosphorylation abundance at all three canonical PDHA1 residues in a dose-dependent manner, with maximal abundance achieved at 5 µM (Fig. 4e). In contrast, in dermal fibroblasts the PDHA1 residue S293 was undetectable while the residues S232 and S300 were detected but not altered by GDC-0339 treatment (Fig. 4e). Critically, the increase in S232, S293, and S300 PDHA1 phosphosites was not due to increased PDHA1 protein levels (Fig. 4f,g). Immunofluorescent imaging showed total PDHA1 levels did not vary in either cardiomyocytes or fibroblasts in hiPSC-cardiac monolayers treated with 5 μM GDC-0339 for 1 hour (Fig. 4f,g). Quantification of the ratio of phospho-PDHA1 (S293) to total PDHA1 showed cardiomyocytes had increased levels of S293 phosphorylation, which was not observed in cardiac fibroblasts (Fig. 4f,h). Furthermore, the PDK inhibitor dichloroacetate (hereafter, PDKi) (Whitehouse et al., 1974), effectively lowered PDHA1 phosphorylation in both cell types (Fig. 4f,h). Western blotting confirmed an increase in phospho-PDHA1 (S293) to total PDHA1 in response to acute GDC-0339 treatment, which was blocked by PDKi (Extended Data Fig. 7). Consistent with inhibition of PDK activity, GDC-0339 treatment over 96 hours resulted in lower oxygen consumption rate compared to DMSO controls (Fig. 4i). These data suggest that GDC-0339 promotes a metabolic switch in cardiomyocytes via activation of PDK signalling.

In summary, we identify compounds that promote proliferation of cardiomyocytes without affecting other cell types. Mechanistically, cardiomyocyte-specific induction of proliferation appears to involve extensive remodelling of the F-actin cytoskeleton and reprogramming to anaerobic metabolism. Furthermore, we identify the PIM kinase inhibitor, GDC-0339, as a tool compound for cardiomyocyte-specific induction of proliferation. Our data reveal that PIM kinase-dependent phosphorylation of PDHA1 is required for shifting cardiomyocytes into a permissive metabolic state for cell division. Our findings suggest that cardiomyocyte-specific cell cycle regulatory mechanisms are potentially druggable, opening future avenues towards pharmacological cardiac repair.

## Online Methods

### Stem cell culture

Ethical approval was obtained by the Murdoch Children’s Research Institute and Human Research Ethics Committee Approval (93025) for the use of human pluripotent stem cells and human embryonic stem cells.

hiPSCs were reprogrammed using Sendai virus from peripheral blood mononuclear cells by the iPSC Derivation and Gene Editing Facility (Murdoch Children’s Research Institute). PB10.5 hiPSCs were generated from cells from a healthy donor (Vlahos et al., 2019). Desmoplakin (DSP) mutant and corrected hiPSC lines were generated as described by Pocock et al. (2025) from cells provided by a patient donor (MCRI Biobank ID; MCHTB11). Embryonic stem cells (ESCs) expressing NKX2-5-^eGFP^ (heterozygous) were generated by Elliott et al. (2011).

Stem cells were maintained at 37 °C and 5 % CO_2_ in mTeSR-Plus on Matrigel-coated tissue culture flasks. Cells were passaged using TrypLE. All cell lines were confirmed as karyotypically normal by the Victorian Clinical Genetics Service and routinely tested for mycoplasma.

### Culture of breast epithelial cells and primary fibroblasts

MCF10A breast epithelial cells and complete culture media were provided by Kristin Brown (Peter MacCallum Cancer Centre). MCF10A cells were maintained in 37 °C and 5 % CO_2_ and passaged using Trypsin.

Primary fibroblasts (Cellosaurus ID: GM21808) were provided by the iPSC Derivation and Gene Editing Facility (Murdoch Children’s Research Institute). Fibroblasts were maintained in 37 °C and 5 % CO_2_ in DMEM high glucose supplemented with 15 % FBS, 1 % Glutamax, 1% NEAA and 1% penicillin–streptomycin. Fibroblasts were passaged using Trypsin.

Generation of a double reporter PB10.5 TNNT2-eGFP PCNA-mScarlet-I hiPSC line by CRISPR/Cas9 gene editing

The TNNT2-eGFP construct was generated by taking a ∼1500bp section from the 3’ end of the human *TNNT2* (cTnT) gene coding sequence and importing it into SnapGene. Left and right targeting arms (approximately 500bp each) were defined. An AscI restriction enzyme site was inserted between the arms, and the adjacent PAM site at the end of the left arm was deleted to allow Cas9 to cut the wild-type allele but not the integrated target allele. This modified ∼1000bp segment was then inserted into a pUC57-kan plasmid backbone. A guide RNA (gRNA; sequences below) was used to direct the Cas9 nuclease (using the pSpCas9n(BB) plasmid) to the *TNNT2* locus.

TNNT2_gRNA_F: 5’ caccgatctttggtgaaggaggcc 3’

TNNT2_gRNA_R: 5’ aaacggcctccttcaccaaagatc 3’

The homology (*TNNT2*-eGFP) and CRISPR/Cas9 with gRNA construct were transfected into wildtype PB10.5 hiPSCs. Transfected cells were isolated via Fluorescence-Activated Cell Sorting (FACS), gating for eGFP-positive/PI-negative cells. The sorted populations were plated onto mouse embryonic fibroblast (MEF) feeders supplemented with ROCK inhibitor. Individual surviving colonies were manually picked, transferred to 48-well plates, and serially expanded into 6-well plates and T25 flasks. Confirmed clones subsequently underwent a secondary single-cell FACS sort into 96-well plates to ensure true clonality before final expansion and adaptation to feeder-free conditions on Matrigel. Final clones were validated by PCR screening to establish zygosity then Sanger sequencing to confirm 100 % sequence fidelity. The final hiPSC clone, with homozygous *TNNT2*-eGFP integration, was validated by differentiation into 2D cardiac monolayers and sarcomere expression of eGFP was confirmed.

The double reporter PB10.5 TNNT2-eGFP PCNA-mScarlet-I hiPSC line was generated using the PCNA-mScarlet-I homology construct (tagging the 3’ end of the *PCNA* gene) and gRNA construct published by Butera et al., (2024). Plasmids were transfected into hiPSCs of the validated PB10.5 TNNT2-eGFP clone. Transfected cells were expanded into Matrigel-coated t75s then directly single cell sorted into Matrigel-coated 96 well plates to generate monoclonal lines. The final clone was selected by microscopy to confirm correct nuclear localisation and appearance of PCNA. The double reporter line was confirmed as karyotypically normal.

All transfection steps were completed by electroporation using the ThermoFisher Neon Transfection System, whereby 1 million hiPSCs were transfected using 2 pulses at 1050 V for 30 ms.

### Stem cell-cardiac differentiation

iPSCs were differentiated into cardiac monolayers according to the protocol described by Voges et al., (2023) using a 15 day protocol. HiPSCs were seeded onto Matrigel-coated flasks four days prior to induction of differentiation (day -4).

iPSC cells were differentiated using a base media of RPMI 1640 Medium with GlutaMAX Supplement, 200 μM AA2P, and 1% penicillin–streptomycin. On days 0, 1 and 2, cells were treated with base media supplemented with 2 % B-27 minus insulin, Activin A (9 ng/ml), BMP-4 (5 ng/ml), FGF-2 (5 ng/ml) and CHIR99021 (1 µM). On day 3, base media was supplemented with 2 % B-27 minus insulin and IWP-4. On day 6, 8 and 10 cells were treated with base media with 2% B-27 including insulin and 5µM IWP-4. The final media change on day 13 used base media containing 2% B-27 including insulin.

### Harvesting day 15 stem cell-cardiac monolayers

hiPSC-cardiac monolayers were washed with Versene solution then detached from tissue culture flasks on differentiation day 15 using pre-warmed Trypsin (3 ml per t25) with 15 min incubation at 37 °C and gentle tapping at 5 min intervals. Trypsin was diluted with 3ml solution of Versene and FCS (1:1). Cells were maintained on ice prior to seeding. Cells were counted and the desired number of cells were centrifuged at 300 g for 4 min then supernatant was removed. Cells were resuspended then seeded in RPMI 1640 Medium with GlutaMAX Supplement, 2% B-27 including insulin, 200 μM AA2P, 1% penicillin–streptomycin, and ROCK inhibitor Y-27632 (10 µM). Cells were seeded in PerkinElmer 384-well PhenoPlates. For the primary screen (fixed), 12,500 cells were seeded in 80 µl per well. For the secondary screen (live imaging), 5,000 cells were seeded in 80 µl per well.

### Flow cytometry of harvested day 15 stem-cardiac monolayers

Samples were stained using the Zombie NIR Fixable Viability Kit (Australian Biosearch) then fixed and stained with Medium A and Medium B from the FIX & PERM Cell Permeabilization Kit (Invitrogen), according to manufacturer protocols. Samples were stained for TNNT2 or ACTN2 to quantify the proportion of cardiomyocytes. Where present, ACTN2-Scarlet intensity was used to validate cardiomyocyte proportion. Flow cytometry was performed using the LSRFortessa X-20.

### Compound screens in hiPSC-cardiac monolayers

24 hr after seeding (differentiation day 16), media was replaced RPMI 1640 Medium with GlutaMAX Supplement, 2% B-27 including insulin, 200 μM AA2P and 1% penicillin–streptomycin without ROCK inhibitor. 48 hr after seeding, media was replaced with ‘maturation media’ previously shown to reduce cell cycle activity (Mills et al., 2017). Maturation media is serum-free medium supplemented with 1 mM glucose with 100 μM palmitate and no insulin (Mills et al., 2017). Cells were maintained in maturation media thereafter, including during compound treatments.

Treatments for compound screens were completed on differentiation day 19. 384-well plates containing compounds pre-dispensed by Compounds Australia were resuspended in 100 µl warmed cell culture media using an automated liquid handling system (Fluent1080, TECAN) to attain a final concentration of 1 µM or 5 µM. The Fluent1080 was used to transfer 80 µl of media containing compound to aspirated cell plates. Four technical replicates were included per compound and dose. Each screening plate contained eight wells with DMSO (negative control) and eight wells with Compound 6.28 (positive control), with four of each at edge wells and four in central wells, to check for edge effects.

### Compound screens in non-cardiac fibroblasts and breast epithelial cells

2000 cells were seeded in 80 µl complete media per well in PerkinElmer 384-well PhenoPlates. The day after seeding, cells were treated in complete culture media for 48 hr prior to fixation. Compounds were prepared and added to cells as described above for hiPSC-cardiac monolayers by resuspending compounds dispensed onto stock plates by Compounds Australia.

### Microscopy imaging of fixed 2D monolayers

HiPSC-cardiac monolayers were imaged using the CellVoyager CV8000 High-Content Screening System (Yokogawa) with a 20x air objective. Five fields of view were imaged per well and maximum intensity projections were generated by CellVoyager software using the three imaging z planes (range 10 µM).

### Image analysis of 2D monolayers

For compound screens, image analysis was performed using CellPathFinder (Yokogawa) using a customised script that segmented non-border nuclei using the Hoechst channel. Mean intensity was measured on a single cell level for all channels. TNNT2 intensity in the segmented nuclear region was used to assign cells as cardiomyocytes or non-cardiomyocytes (cardiac fibroblasts).

To quantify actin morphology, whole images were analysed in CellProfiler (Carpenter et al., 2006). The module ‘MeasureImageIntensity’ was used to calculate intensity. The module ‘MeasureTexture’ was used to calculate Haralick features: Variance, Contrast, Correlation, Angular Second Moment (ASM), and Entropy (Haralick et al., 1973). A horizontal (0 °) orientation and scale factor of 3 was used for quantification.

### Stem cell-cardiac organoid generation and culture

Heart-Dyno inserts were manufactured using the SYLGARD 184 Silicone Kit (PDMS) at a 10:1 ratio of polymeric base to the crosslinking curing agent. Instead of clear tissue culture plates as previously described (Voges et al., 2023), Heart-Dyno inserts were attached onto a 96-well PerkinElmer PhenoPlate, which has black well walls and a plastic bottom with high optical quality, enabling direct imaging on this plate without mounting onto microscopy slides.

Stem cell-cardiac organoids were generated according to the protocol described in (Voges et al., 2023) with the updated adaptations from Pocock et al., (2025). Updates include the use of CHIR99021 (2 µM) in the organoid matrix mixture on day 15, and addition of 3 μM DY131 (ERR agonist) and 10 μM MK8722 (AMPK activator) on days 25 and 28.

### Compound treatments of stem cell-cardiac organoids

Treatments were performed on day 29 of the differentiation protocol in weaning media or, where specified, day 22 in maturation media. All stem-cardiac organoids were fixed 48 hr after compound addition. GDC-0339 was sourced from MedChemExpress (#HY-16976) and GSK3i-IX was sourced from Sigma-Aldrich (#361550).

### Force analysis of stem cell-cardiac organoids

Stem cell-cardiac organoids were imaged using the Leica Thunder microscope in an environmental chamber set to 37 °C and 5% CO_2_. Force analysis was performed using the protocol described by Mills et al., (2017), whereby images were taken of each organoid for 10 seconds at 50 Hz. Contractility analysis was performed using Tempo.ai software (Dynomics).

### Immunofluorescence staining of 2D monolayers

Stem cell-derived cardiac monolayers, non-cardiac fibroblasts and MCF10A breast epithelial cells were fixed with paraformaldehyde at a final concentration of 2-4 % for 1 hr at RT. Following fixation, samples were washed with PBS three times then permeabilised with 0.2 % TritonX-100/PBS for 30 min at RT. Following PBS washes, samples were blocked in 2 % BSA/PBS for at least 1 hr at RT or overnight at 4 °C. Primary antibodies were diluted in 2 % BSA/PBS added overnight at 4 °C. Samples were incubated with Hoechst (1:1000) with secondary antibodies diluted in 2 % BSA/PBS for 90 min at RT. Samples were washed three times with PBS and stored at 4 °C then acclimated to RT for imaging.

### Live imaging of hiPSC-cardiac monolayers

Live imaging was completed using the CV8000 spinning disk confocal system (Yokogawa) with an environmental chamber set to 37 °C and 5 % CO_2_. Imaging was commenced 30 min after compound addition to allow for plate acclimation to 37 °C. Images were taken at 20 min intervals for 145 timepoints at a single z position.

### Cardiocycle.py analysis of live imaged 2D hiPSC-cardiac monolayers

Live imaged 2D hiPSC-cardiac monolayers expressing PCNA-mScarlet-I and TNNT2-eGFP were analysed using our custom pipeline Cardiocycle.py (DOI: 10.5281/zenodo.20423193). Cardiocycle.py implements three key processing steps: 1) restructuring and resizing of image files, 2) nucleus segmentation based on PCNA labelling, 3) single cell tracking and stratification of tracks by TNNT2 status. In the first step, image files are reorganised from a flat to modular structure and renamed to include essential metadata (experiment date and compound dose) while removing redundant information. Images are also resized to 700 x 700 px, which was optimised as sufficient resolution for accurate tracking while minimising image file sizes to speed up downstream image processing. PCNA-labelled nuclei were segmented using the machine learning module in Cellpose (Stringer and Pachitariu, 2025). Supervised learning was performed by manually providing masks for over 1000 non-border nuclei across 30 images. Training images were used from six biological experiments derived from independent hiPSC-cardiac differentiations to maximise variation in PCNA measurements and nuclear morphology. Models were iteratively trained with 200 epochs per model, finalised at 30 models at which nuclei segmentation was visually assessed as accurate. To perform single cell tracking, a script was customised in CellProfiler (Carpenter et al., 2006) to perform tracking using segmented output from Cellpose. The entire workflow is compiled into a standalone script that can be submitted to a high-performance computing (HPC) system.

### Immunofluorescence staining of stem cell-cardiac organoids

Stem cell-derived human cardiac organoids were fixed and stained directly in 96-well PerkinElmer PhenoPlates. Fixation was performed at RT for 1 hr with 30 µl 16% PFA added to the existing cell culture media (150 µl). After three PBS washes, organoids were permeabilised for 1 hr with 0.25 % TritonX-100 and 5 % FBS in PBS. After a PBS wash, organoids were incubated at 4 °C with a blocking solution (2 % BSA and 5 % goat serum in PBS) for at least 1 hr. Antibody staining was completed with the same blocking solution at 4 °C on a rocker. Primary antibodies were added overnight, washed 3 times with PBS with 1 hr incubations, then secondary antibodies were added with Hoechst (1:1000). After 3 times 1 hr washes with PBS, organoids were cleared using 60 % glycerol and 2.5 M fructose in dH_2_O for 30 min. Clearing solution was diluted twice with 200 µl PBS per well prior to imaging.

### Microscopy imaging of fixed stem cell-cardiac organoids

Human iPSC-cardiac organoids were imaged using a 20X air objective (NA 0.8, resolution 3.3 px/µm) with a 40 µm pinhole on the Andor Dragonfly 200 Spinning Disk Confocal Microscope. Each organoid was imaged with three horizontally tiled fields (10 % overlap). Organoids were imaged over a range of 200 µM with 9 z-slices.

### Image analysis of fixed stem cell-cardiac organoids

Image fields were stitched using Fusion software and exported as .ims files. ims files were converted to .tiff files and maximum intensity projections were generated then separated by channel using FIJI (ImageJ). Nuclei were segmented using the Hoechst channel or NKX2-5 (reporter or antibody staining) where present. Segmentation was performed using the ImageJ StarDist plugin. Downstream image analysis was performed using CellProfiler, importing segmented nuclei as primary objects then measuring channel intensities in the nuclear regions of non-border nuclei.

### Sample preparation for phosphoproteomics

Day 15 hiPSC-cardiac monolayers were harvested and seeded in Matrigel-coated 6-well plates with 1.4 million cells per well in RPMI 1640 Medium with GlutaMAX Supplement, 2% B-27 including insulin, 200 μM AA2P and 1% penicillin–streptomycin with ROCK inhibitor and changed the following day to the same media without ROCK inhibitor. On day 16 and 17, media was changed to maturation media. All media changes and compound treatments used 2 ml media per well. Compound treatment and processing was performed on day 18.

For fibroblasts, 320,000 cells were seeded per well on uncoated TC-treated 6-well plates in 2ml RPMI-glutamax supplemented with 15 % FBS, 1% NEAA and 1 % penicillin–streptomycin. The following day, samples were treated in serum-free RPMI-glutamax with 1% NEAA and 1 % penicillin–streptomycin.

For treatment and processing for phosphoproteomics, culture media was aspirated and replaced with 2 ml media with compound (GDC-0339, GSK3i-IX, C6.28 or DMSO) made by serial dilutions. Samples were processed in batches of 28 wells (7 treatments x 4 biological replicates) to maintain sample quality. After 1 hr compound treatment in a 37 °C incubator, plates were placed on ice. Media was then aspirated and cells were washed three times with cold TBS. 150 µl lysis buffer (prepared the same day) was then added per well. Lysis buffer consisted of 4 % Sodium deoxycholate and 1 M Tris (pH 8.5) in Milli-Q ultrapure water. After lysis buffer was added to all 28 wells, samples were scraped then pipette into a pre-chilled 1.5 ml Eppendorf then transferred to a hot block pre-heated to 95 °C for 5 min then returned to ice. Each sample was sonicated for 5 seconds three times then returned to ice. 10 µl aliquots per sample were used to estimate protein concentrations using the Pierce BCA assay kit (Thermo Fisher Scientific, #23225). Samples were stored at -80 °C prior to further processing.

### LC-MS/MS measurement

Enriched phosphopeptides in MS loading buffer (2 % ACN, 0.3 % TFA) were loaded onto in-house fabricated 55 cm columns with a 75 µm I.D. and packed with 1.9 µm C18 ReproSil Pur AQ particles using a Vanquish Neo HPLC coupled to an Orbitrap Astral mass spectrometer (Thermo Fisher Scientific). Column temperature was maintained at 60 °C in a Sonation column oven, and peptides separated using a binary buffer system comprising 0.1 % formic acid (buffer A) and 80 % ACN plus 0.1 % formic acid (buffer B), at a flow rate of 400 nl min–1 with a gradient of 3-19 % buffer B over 20 min followed by 19-41 % buffer B over 10 min, resulting in ∼ 30 min gradients. Peptides were analysed with one full scan (380-980 m/z, R = 240,000) at a target of 5 × 10^6^ ions, followed by 300 data-independent acquisition (DIA) MS/MS scans (380-980 m/z) with HCD (target 5 x 10^4^ ions, maximum injection time 3.5 ms, isolation window 2 m/z, NCE 24 %), with fragments detected in the Astral.

### Phosphoproteome data processing and analysis

Raw data were analysed using direct DIA analysis in the Spectronaut software (version 19.3.241023.62635). A spectral library was generated against the Human UniProt FASTA database (accessed in March 2025). Default settings were used for search parameters. The following preprocessing and analyses were performed in the R software environment version 4.4.2. To remove unreliable phosphosites, we first removed all phosphosites that had a localisation probability of < 0.5, then any remaining phosphosites with a maximum localisation probability < 0.75 were further removed (retaining only class I phosphosites). We further filtered out phosphosites missing in at least 1/3^rd^ of hiPSC-cardiomyocyte samples and all values < 3 were converted to missing values as they were considered unreliable measurements indistinguishable from background noise. Raw intensities were log_2_ transformed, after which we performed centre-median normalisation.

The Likelihood Ratio Test (LRT) in the DESeq2 package in R was used to identify differentially regulated regulated phosphosites across the 7-point dose curve for each compound (GDC-0339, GSK3i-IX and C6.28) independently. Significant phosphosites were filtered based on a log2foldchange > 1 and padj < 0.05. The same test was performed for hiPSC-cardiac cells and fibroblasts independently. Differentially regulated phosphosites identified in hiPSC-cardiac cells but also differentially regulated in fibroblasts were filtered out to attain cardiomyocyte-specific phosphosites. To identify core signalling pathways across compounds, cardiomyocyte-specific phosphosites present in at least two of three compounds, and regulated in the same direction (i.e. all upregulated or all downregulated) were filtered for. Kinase-Substrate Enrichment Analysis (KSEA) was performed using KSEAapp (Wiredja et al., 2017) and Gene Set Enrichment Analysis (GSEA) was performed using clusterProfiler (Yu et al., 2012).

### Western blotting

Human iPSC-cardiac monolayers treated with compounds for 48 hr were lysed using RIPA buffer. Lysates were clarified by centrifugation (20,000 × g, 15 min), and protein concentration was determined using the Pierce BCA Protein Assay Kit (Thermo Fisher, #23225). Samples were mixed with 1X Laemmli buffer and denatured by heating to 70 °C for 10 min prior to loading. 10 μg protein (15 μL total volume) was loaded per lane and resolved on a 4-20 % Mini-PROTEAN TGX precast gel (Bio-Rad, 4561096EDU) at 70 V for 20 min, then increased to 115 V for 60 min, in 1X Tris/Glycine/SDS running buffer (Bio-Rad, 1610772EDU). The Precision Plus Protein Dual Colour Standard (Bio-Rad, #1610374) was run alongside samples.

Proteins were transferred to a PVDF membrane (Immun-Blot PVDF Membrane, 1620177, Bio-Rad) at 0.3 A for 2 hr at 4 °C in 1X Tris/Glycine transfer buffer (Bio-Rad, 1610771EDU) supplemented with 10% methanol. Following transfer, membranes were briefly rinsed in 1X TBS-T, stained with Ponceau S (P7170-1L, Sigma-Aldrich) to visualise total protein, and imaged on a Bio-Rad GelDoc system (12009077). Residual Ponceau S stain was removed by washing in 1X TBS-T (2 min), and membranes were blocked in 5 % milk in TBS-T for 30 min at room temperature.

Primary antibodies (anti-PDHA1 (1:2000; Abcam, ab110330) and anti-phospho-PDHA1 (Ser293) (1:1000; Abcam, ab92696)) were diluted in 5% milk in TBS-T and incubated with membranes for 36 hr at 4 °C with constant rolling. Membranes were then washed (3x 15 min) in 1X TBS-T and incubated with HRP-linked secondary antibody (anti-mouse IgG, CST #7076, or anti-rabbit IgG, CST #7074; 1:5000 in 5% milk in TBS-T) for 1 hr at room temperature with constant rolling, in the dark. Membranes were washed (3 x 15 min) in 1X TBS-T, then incubated with Clarity ECL substrate (Bio-Rad, #1705060) for 2 min, and protein bands were visualised using an Amersham Imager 680. Band densitometry was performed in ImageJ.

### Oxygen consumption rate assay

Live-cell respiration was measured in real-time using Resipher (Lucid Scientific). PB10.5 hiPSC-cardiac monolayers were cultured in matrigel-coated tissue culture 96-well plates. On day 19, cultures were treated with GDC-0339 at 5 µM or DMSO control. The Resipher sensor lid was transferred to the culture plate and media O_2_ was measured with an operating height between 1,600 and 2,100 µm. Respiration rates of GDC-0339-treated hiPSC-cardiac cells were measured as pmol/min and normalised to untreated DMSO-treated wells on the same plate.

### Statistical analysis

Data is presented as n = biological/experimental replicates, unless otherwise stated. Statistical tests were performed in R using the rstatix package and displayed using the ggplot2 and ggpubr packages. Benjamini-Hochberg adjustment was applied to all statistical tests between more than two groups. Values of p < 0.05 (or padj where relevant) were considered statistically significant.

## Data availability

All proteomics data have been deposited on PRIDE (ID: PXD078756) and will be publicly available following publication of this manuscript. Cardiocycle.py has been deposited on Github via Zenodo and is readily available (DOI: 10.5281/zenodo.20423193). Further information and requests for resources and reagents should be directed to and will be fulfilled by the lead contacts.

## Acknowledgements

We thank Compounds Australia for the provision of compound libraries and technical support with plate design.

## Author Contributions

F.B., K.I.W., D.E. and E.P. conceptualised the study. F.B., R.M., I.C. L.H.H.L., J.L, S.R.H. S.B.S. and T.C. performed experiments. F.B., B.H. and H.H. analysed data.

K.B., E.K., T.M., A.Z., E.B.K. and J.M. provided cells and reagents. A.H.G. and S.J.H. supported high-throughput screening and phosphoproteomics respectively. E.P and D.E obtained funding and supervised the project. F.B. drafted the manuscript. E.P. and D.E. edited the manuscript. All authors approved the manuscript for submission.

---

**Extended Data Fig. 1.**
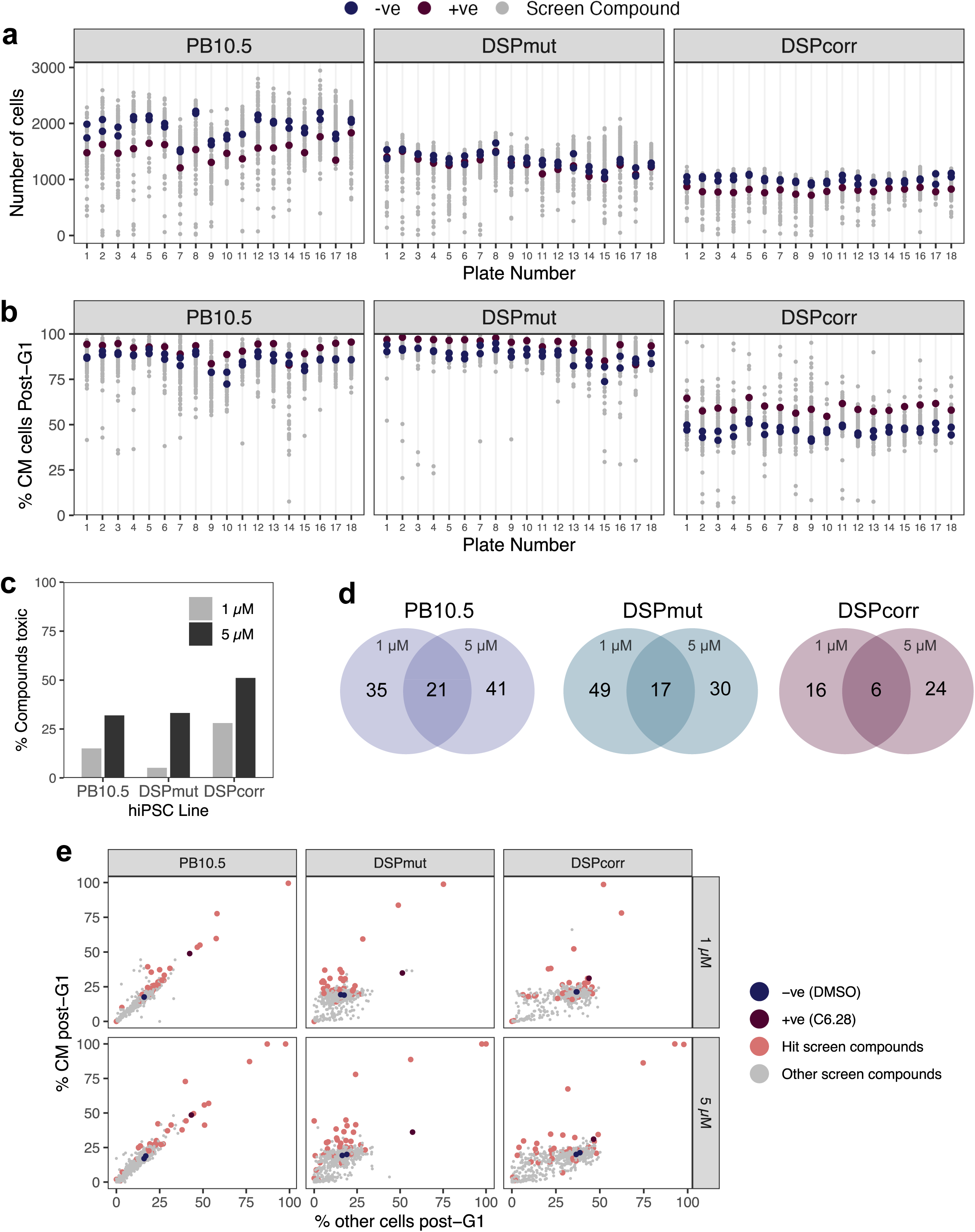
Quality controls and hit selection from fixed hiPSC-cardiac monolayers for cardiomyocyte proliferative compounds to validate with Cardiocycle. a. Mean number of hiPSC-cardiac cells per well in primary screen by compound and hiPSC line. Each point is a screen compound (total 775). The negative (-ve) control is DMSO and positive (+ve) control is 5 µM Compound 6.28. b. Mean percentage of cardiomyocytes (CM) positive for the post-G1 marker pRB by screen compound and hiPSC line. c. Percentage of compounds categorised as toxic by cell line and compound dose. Toxicity was measured as a significant reduction in cardiomyocyte frequency compared to the DMSO control. d. Number of proliferative compound hits (z-score > 1.75) by cell line and compound dose. Final hits (39 compounds) were selected as hits in two or more cell lines irrespective of compound dose. e. Percentage of cardiomyocytes (TNNT2+ cells) versus non-cardiomyocytes (TNNT2-cells) in fixed hiPSC-cardiac monolayers positive for the post-G1 cell cycle marker phospho-RB following treatment with 775 compounds at 1 or 5 µM. 39 compounds (orange) were followed up with the Cardiocycle pipeline.

**Extended Data Fig. 2.**
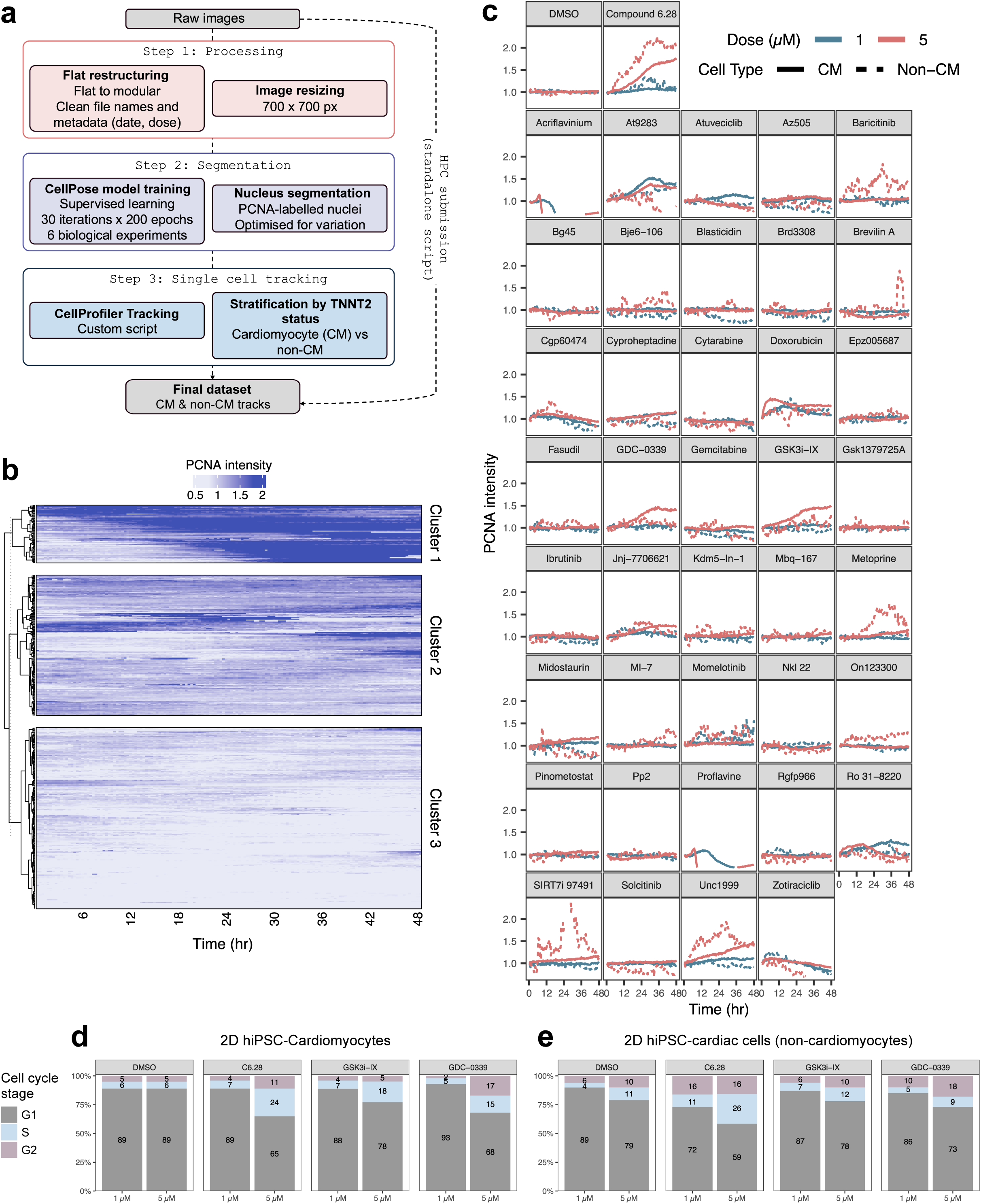
Cardiocycle identifies clusters of hiPSC-cardiomyocyte cell cycle activity and assesses the cardiomyocyte-specificity of compounds. a. Schematic overview of Cardiocycle pipeline. HiPSCs expressing PCNA-mScarlet-I and TNNT2-eGFP are differentiated into hiPSC-cardiac monolayer, enabling stratification of hiPSC-cardiac cells by cell cycle state and cell type (cardiomyocyte/CM or non-cardiomyocyte/non-CM). HiPSC-cardiac monolayers are imaged by timelapse microscopy for 48 hr with 20 min intervals. Here, we assessed 39 compounds in addition to a negative (DMSO) and positive (Compound 6.28) control. Raw microscopy images are automatically organised into a modular structure and renamed with input (user selected) metadata, such as date and dose. The Cellpose training model (supervised machine learning) was used with images of PCNA-mScarlet-I from 6 biological replicates to build a model that segments PCNA-labelled nuclei. Single cell tracking is performed using CellProfiler, inputting segmented nuclei, PCNA-mScarlet-I and TNNT2-eGFP images matched by field and timepoint. Tracked nuclei are then stratified by TNNT2 status (positive = cardiomyocyte). Data for each compound are then analysed as averages by cell type and treatment (e.g. dose) or as single nucleus tracks. b. Clustered heatmap of single nucleus tracks of PCNA intensity over 48 hr for hiPSC-cardiomyocytes (TNNT2+ cells) treated with DMSO, Compound 6.28, GSK3i-IX or GDC-0339. c. Tracks of mean PCNA-mScarlet-I intensity over time for DMSO (negative control), and Compound 6.28 (positive control), and the 39 hit compounds from fixed hiPSC-cardiac monolayers, Tracks are stratified by cell type and dose. PCNA-mScarlet-I intensity was quantified by living imaging of hiPSC-cardiomyocyte monolayers at 20 min intervals. Cardiomyocyte/TNNT2+ cells are represented by a solid line and non-cardiomyocyte/TNNT2-cells are represented by a dashed line. d. Proportion of hiPSC-cardiomyocytes in each cell cycle stage (G1, S or G2) classified using PCNA appearance, showing an increase in S and G2 nuclei by Compound 6.28 (C6.28), GSK3i-IX and GDC-0339. e. Proportion of non-cardiomyocytes (hiPSC-cardiac fibroblasts) in hiPSC-cardiac monolayers in each cell cycle stage (G1, S or G2) classified using PCNA appearance, showing an increase in S and G2 nuclei by Compound 6.28 (non-specific proliferation) and limited change by GSK3i-IX and GDC-0339 (cardiomyocyte-specific proliferation).

**Extended Data Fig. 3.**
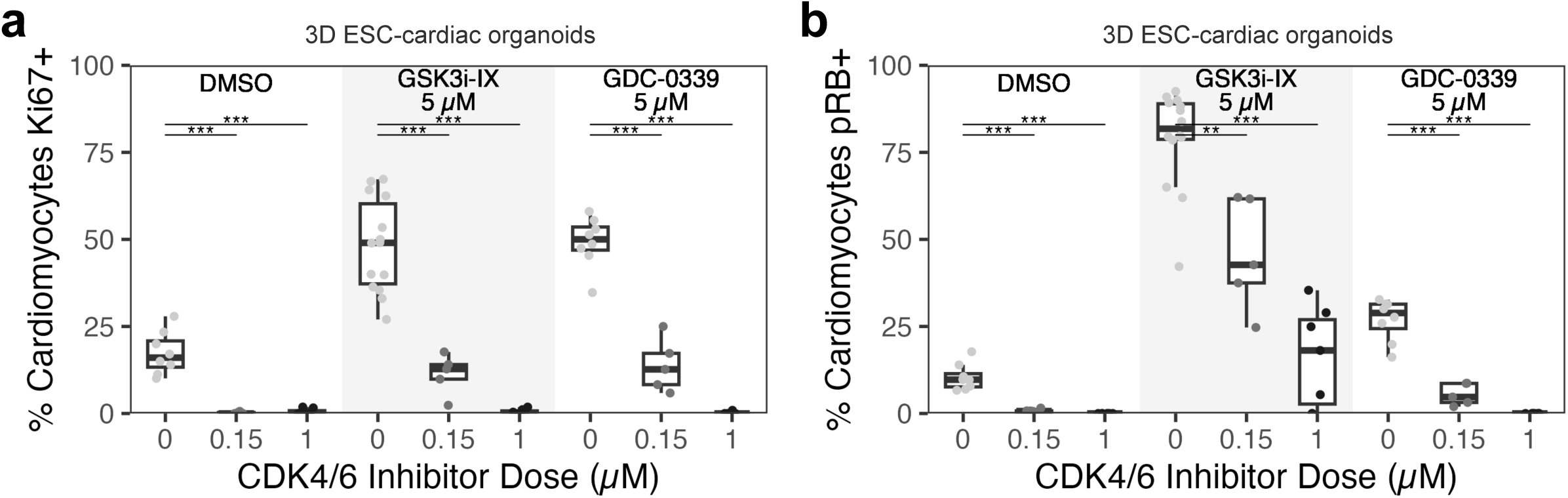
GSK3i-IX and GDC-0339 induce cardiomyocyte proliferation reversed by CDK4/6 inhibition in ESC-cardiac organoids. a. Percentage of cardiomyocytes (NKX2-5-GFP+ cells) in ESC-cardiac organoids positive for the proliferation marker Ki67. Cells were treated in weaning media with DMSO, GSK3i-IX (5 µM) or GDC-0339 (5 µM) and co-treated with the CDK4/6 inhibitor Palbociclib (at 0, 0.15 or 1 µM) for 48 hr from day 28-30 of the differentiation protocol. n = 5-7 organoids per condition (1 experiment). b. Percentage of cardiomyocytes (NKX2-5-GFP+ cells) in ESC-cardiac organoids positive for the proliferation marker phospho-RB. Cells were treated in weaning media with DMSO, GSK3i-IX (5 µM) or GDC-0339 (5 µM) and co-treated with the CDK4/6 inhibitor Palbociclib (at 0, 0.15 or 1 µM) for 48 hr from day 28-30 of the differentiation protocol. n = 5-7 organoids per condition (1 experiment).

**Extended Data Fig. 4.**
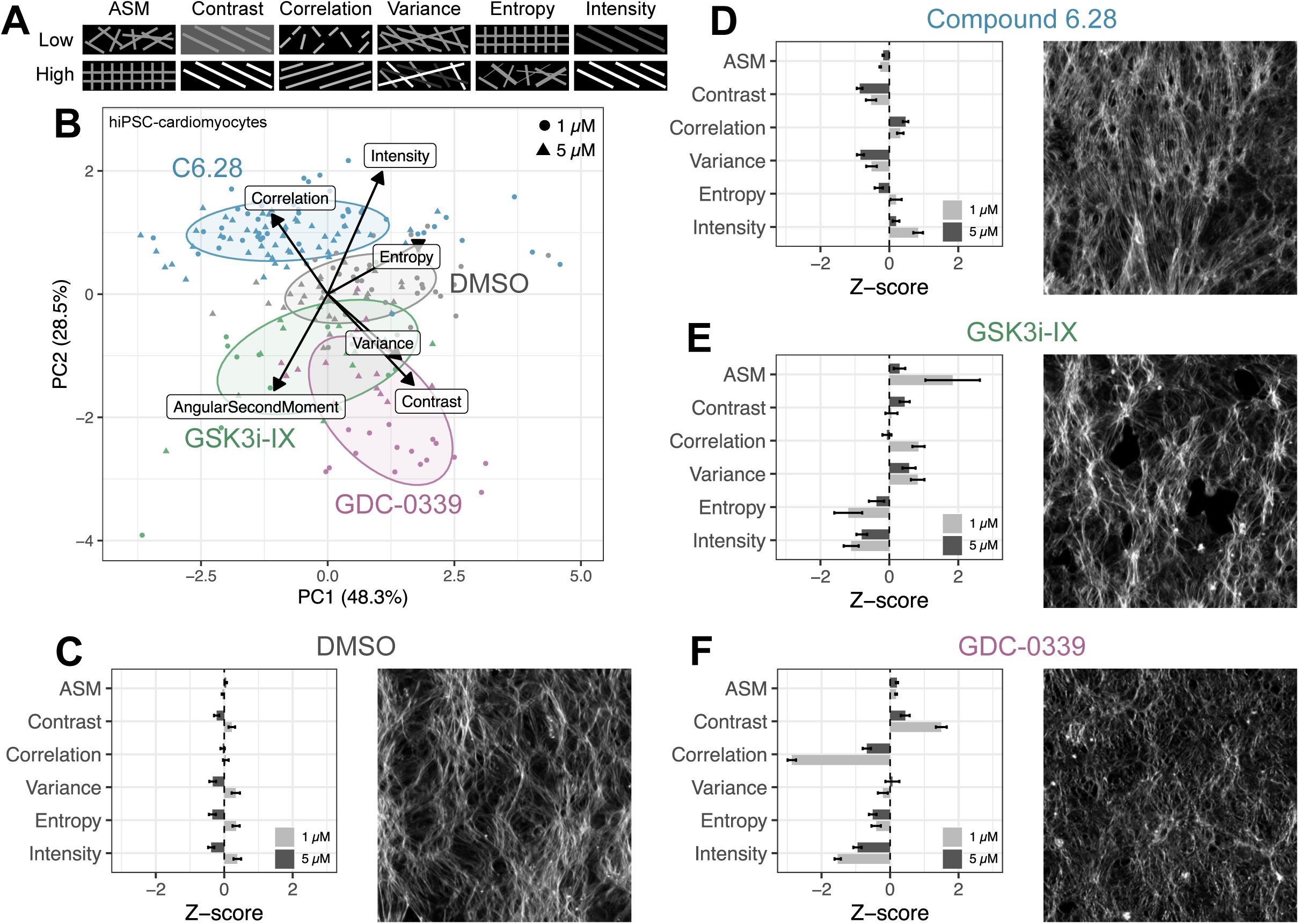
Quantification of F-actin intensity and morphology in hiPSC-cardiac monolayers. a. Filamentous (F)-actin intensity and Haralick texture features. Intensity measures raw F-actin abundance, contrast is intensity relative to surrounding pixels, and variance measures heterogeneity in intensity across the field of view. Correlation quantifies dependency of pixel intensity on neighbouring pixels, with long or parallel filaments having high correlation while short or offset filaments having low correlation. Angular Second Moment (ASM) and entropy measure regularity/uniformity of F-actin across the field of view and are generally inversely correlated. b. PCA of F-actin features quantified in hiPSC-cardiac monolayers treated with DMSO, Compound 6.28, GSK3i-IX or GDC-0339 (1 and 5 µM) for 48 hr and stained with phalloidin. Each point represents a technical replicate. c-f. Quantification of F-Actin intensity and morphology features and representative microscopy images of F-actin (from phalloidin staining) for hi-PSC cardiac monolayers. Cells were treated 48 hr with DMSO, Compound 6.28, GSK3i-IX or GDC-0339 at 1 and 5 µM. Measurements are z-score normalised to the DMSO control by feature.

**Extended Data Fig. 5.**
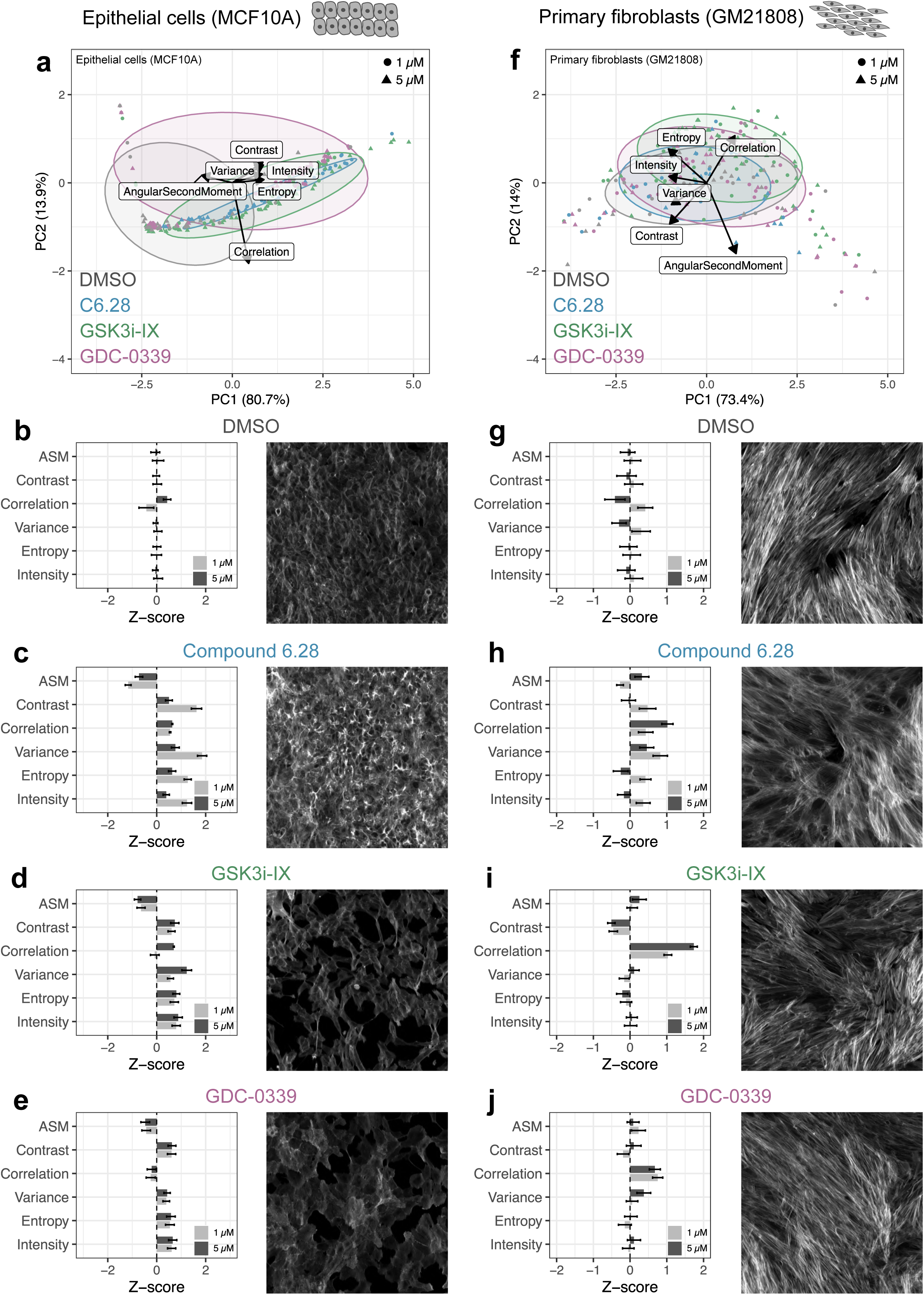
Quantification of F-actin intensity and morphology in MCF10A epithelial cells and GM21808 fibroblasts. a. PCA of F-actin features quantified in MCF10A breast epithelial cells and treated with DMSO, Compound 6.28, GSK3i-IX or GDC-0339 (1 and 5 µM) for 48 hr and stained with phalloidin. Each point represents a technical replicate. b-e. Quantification of F-Actin intensity and morphology features and representative microscopy images of F-actin (from phalloidin staining) for MCF10A breast epithelial cells following 48 hr compound treatment. Measurements are z-score normalised to the DMSO control by feature. f. PCA of F-actin features quantified in GM21808 primary fibroblasts and treated with DMSO, Compound 6.28, GSK3i-IX or GDC-0339 (1 and 5 µM) for 48 hr and stained with phalloidin. Each point represents a technical replicate. g-j. Quantification of F-Actin intensity and morphology features and representative microscopy images of F-actin (from phalloidin staining) for GM21808 primary fibroblasts following 48 hr compound treatment. Measurements are z-score normalised to the DMSO control by feature.

**Extended Data Fig. 6.**
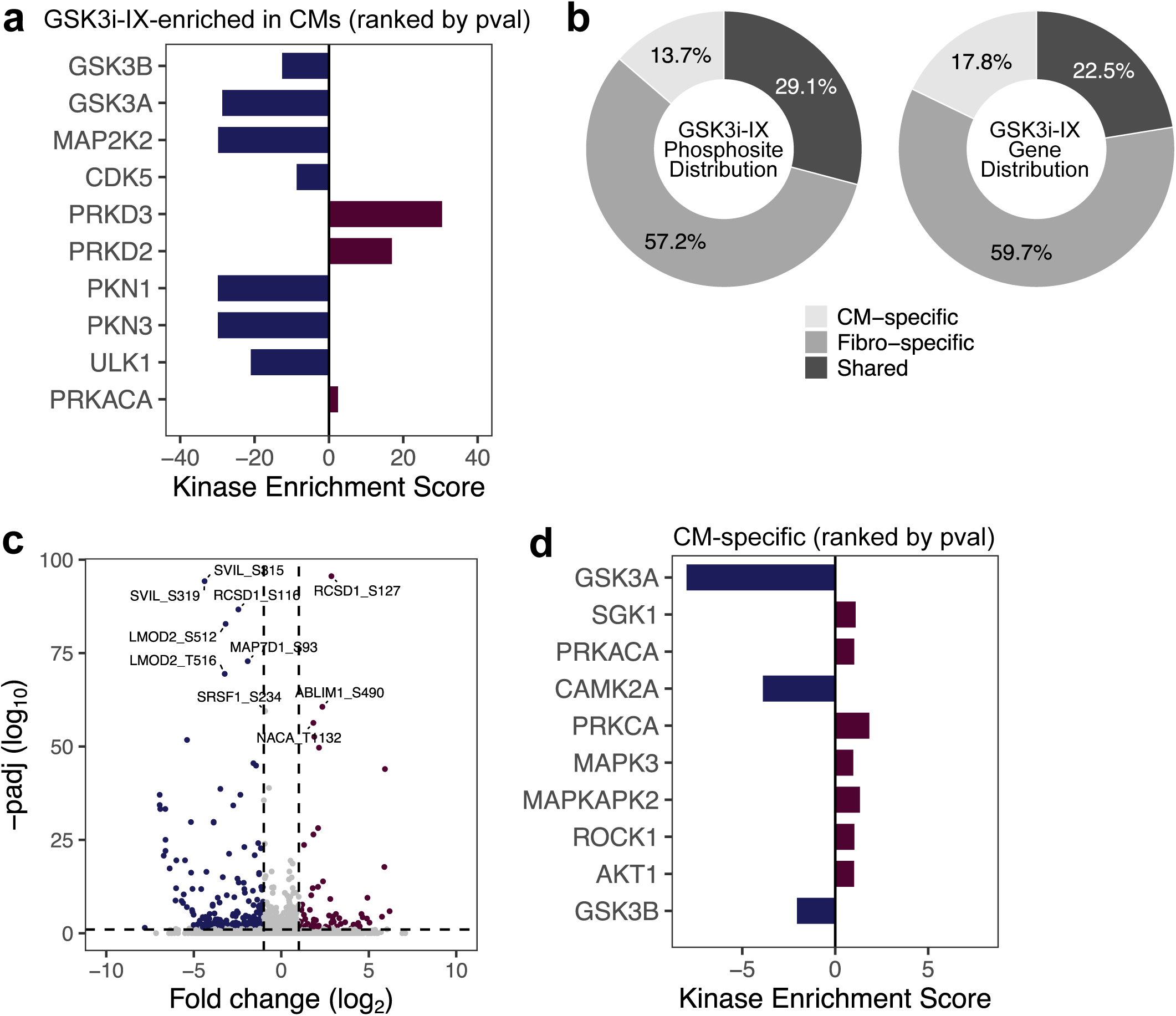
GSK3i-IX displays cell type-specific phosphosignatures. a. Kinases identified by KSEA for differentially-regulated phosphosites by GSK3i-IX in hiPSC-cardiac cells (irrespective of effect in dermal fibroblasts). Upregulated kinases (positive enrichment score) are in red, downregulated kinases (negative enrichment score) are blue. b. Left: percentage of differentially regulated phosphosites by GSK3i-IX classed as statistically significant (DESeq2 Likelihood Ratio Test padj < 0.05, |log2foldchange| > 1) for cardiomyocytes alone (CM-specific), fibroblasts alone (Fibro-specific), or in both cell types (shared). Right: percentage of cardiomyocyte-specific, fibroblast-specific, or shared genes associated with the differentially regulated phosphosites by GSK3i-IX. c. Volcano plot of statistical significance against log2 fold-change of cardiomyocyte-specific phosphosites following GSK3i-IX treatment (7-point dose curve; DESeq2 Likelihood Ratio Test padj < 0.05, |log2foldchange| > 1). The top 10 differentially regulated phosphosites are labelled by protein name and residue. Upregulated phosphosites (padj < 0.05, log2foldchange > 1) are in red, downregulated phosphosites (padj < 0.05, log2foldchange < -1) are in blue. d. Kinases identified by KSEA for GSK3i-IX-regulated cardiomyocyte-specific phosphosites. Upregulated kinases (positive enrichment score) are in red, downregulated kinases (negative enrichment score) are blue.

**Extended Data Fig. 7.**
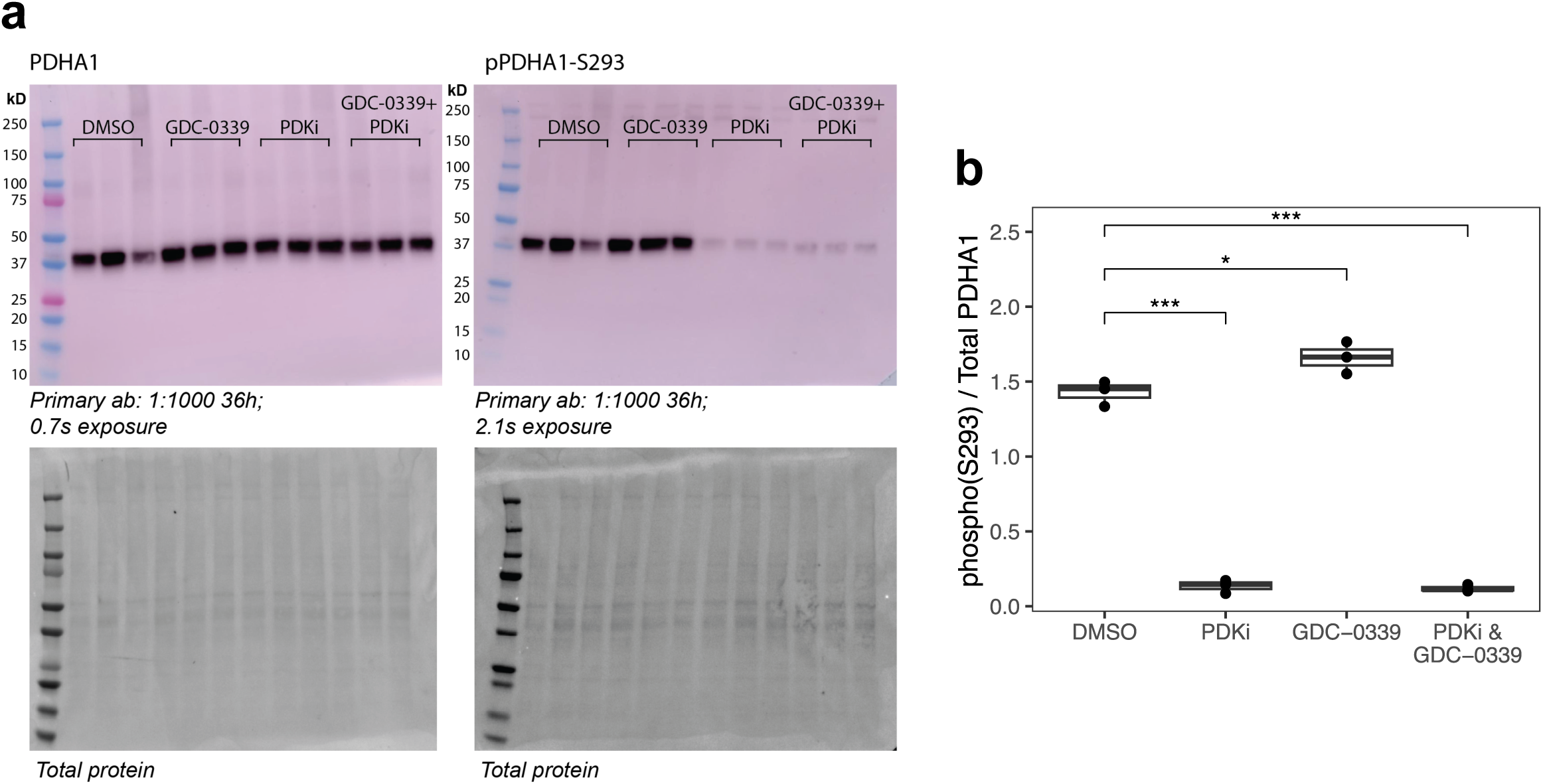
Western blot analysis of phospho(S293)-PDHA1 versus total PDHA1 in hiPSC-cardiac monolayers. a. Western blots showing phosphorylated PDHA1 (S293) and total PDHA1 levels in DMSO, PDKi (10 mM), GDC-0339 (5 µM) or combination of PDKi (10 mM) and GDC-0339 (5 µM). n = 3 replicates per condition. Ponceau staining was used as a protein loading control. b. Quantification of phosphorylated PDHA1 (S293) relative to total PDHA1 from Western blots in (a). n = 3 replicates per condition.

